# The CPEB3 ribozyme modulates hippocampal-dependent memory

**DOI:** 10.1101/2021.01.23.426448

**Authors:** Claire C. Chen, Joseph Han, Carlene A. Chinn, Xiang Li, Mehran Nikan, Marie Myszka, Liqi Tong, Timothy W. Bredy, Marcelo A. Wood, Andrej Lupták

**Author notes:** Correspondence to: Andrej Lupták. Department of Pharmaceutical Sciences, University of California–Irvine, Irvine, California 92697, United States. Marcelo A. Wood. Department of Neurobiology and Behavior, Center for the Neurobiology of Learning and Memory, University of California–Irvine, Irvine, California 92697, United States. Cognitive Neuroepigenetics Laboratory, Queensland Brain Institute, The University of Queensland, Brisbane, QLD 4072, Australia.

## Abstract

A self-cleaving ribozyme mapping to an intron of the cytoplasmic polyadenylation element binding protein 3 (*CPEB3*) gene has been suggested to play a role in human episodic memory, but the underlying mechanisms mediating this effect are not known. The ribozyme’s self-scission half-life matches the time it takes an RNA polymerase to reach the immediate downstream exon, suggesting that the ribozyme-dependent intron cleavage is tuned to co-transcriptional splicing of the *CPEB3* mRNA. Here we report that the murine ribozyme modulates its own host mRNA maturation in both cultured cortical neurons and the hippocampus. Inhibition of the ribozyme using an antisense oligonucleotide leads to increased CPEB3 protein expression, which enhances polyadenylation and translation of localized plasticity-related target mRNAs, and subsequently strengthens hippocampal-dependent long-term memory. These findings reveal a previously unknown role for self-cleaving ribozyme activity in regulating experience-induced co-transcriptional and local translational processes required for learning and memory.

## Introduction

Self-cleaving ribozymes are catalytic RNAs that accelerate site-specific scission of their backbone [1]. Several mammalian ribozymes have been identified [2-9], including a functionally-conserved sequence that maps to the second intron of the cytoplasmic polyadenylation element binding protein (*CPEB3*) gene [4, 10, 11] (Fig 1A). The CPEB3 ribozyme shares its secondary structure and catalytic mechanism with the hepatitis delta virus (HDV) ribozymes [10, 12]. HDV-like ribozymes are widespread among genomes of eukaryotes [13-16] and their biological roles include processing of rolling-circle transcripts during HDV replication [2, 3], 5′-cleavage of retrotransposons [14-16], and in one bacterial example, potentially metabolite-dependent regulation of gene expression [17]. However, the function of the mammalian ribozyme is unknown. In humans, a single nucleotide polymorphism (SNP) at the ribozyme cleavage site leads to a 3-fold higher rate of *in vitro* self-scission, which correlates with poorer performance in an episodic memory task [4, 18] and suggests that the ribozyme activity might play a role in memory formation.

**Fig 1.**
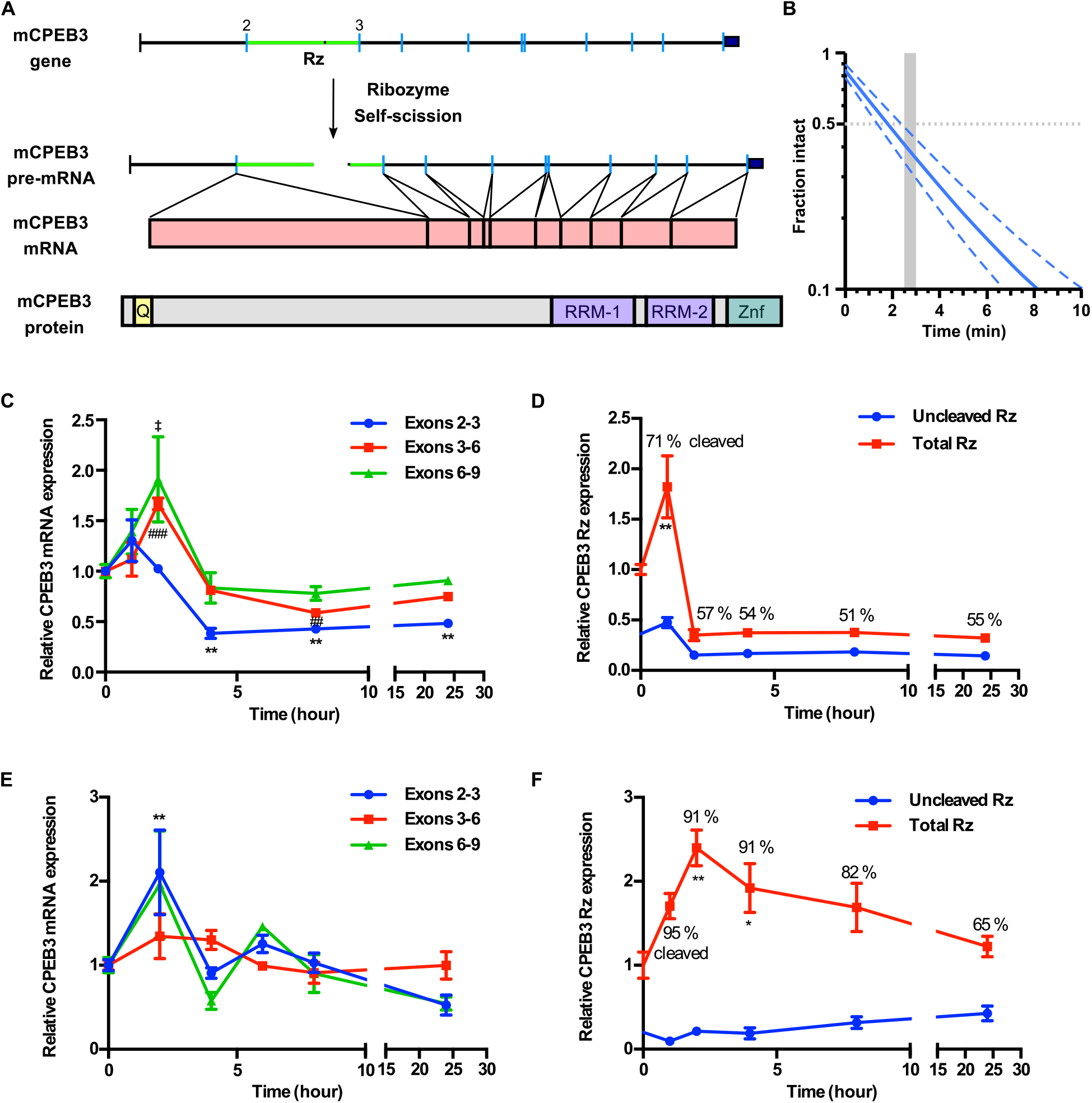
CPEB3 ribozyme activity and its effect on primary cortical neurons. (A) Schematic representation of mouse *CPEB3* gene and its products. Rz denotes the ribozyme location in the 2^nd^ intron (green) between the 2^nd^ and 3^rd^ exons. Co-transcriptional self-scission is shown by a break in the pre-mRNA 2^nd^ intron. Fully-spliced mRNA is shown independent of the ribozyme activity. (B) Co-transcriptional self-cleavage activity of a 470-nt construct, incorporating the 72-nt ribozyme, which cuts the transcript 233 nts from the 5′ terminus (see Table S1 for kinetic parameters of this and other constructs). Log-linear graph of self-cleavage is shown with solid blue line (dashed lines show ± standard deviation). Gray dotted line indicates mid-point of self-cleavage (with resulting t_1/2_ of ∼2 min). Gray bar indicates the approximate time range for RNAPII to reach from the ribozyme to the 3^rd^ exon, at which point ∼40% of the intron would remain intact. (C) KCl stimulation profile of the *CPEB3* gene showing induction of spliced CPEB3 exons (one-way ANOVA with Sidak’s *post hoc* tests. \**P* < 0.05, \*\**P* < 0.01, ##*P* < 0.01, ###*P* < 0.001, ‡*P* < 0.05). (D) KCl stimulation profile of CPEB3 ribozyme expression (uncleaved and total). Cleaved ribozyme fraction is calculated as [(total ribozyme – uncleaved ribozyme)/total ribozyme] and shown as % cleaved. (E) Expression of CPEB3 mRNA exons 2–3 is upregulated 2 hours after glutamate stimulation (one-way ANOVA with Sidak’s multiple comparisons *post hoc* test. \*\**P* < 0.01). (F) Glutamate stimulation induces an increase in CPEB3 ribozyme levels at 2-hour time point (one-way ANOVA with Sidak’s multiple comparisons *post hoc* test. \**P* < 0.05, \*\**P* < 0.01). Data are presented as mean ± SEM.

CPEBs are RNA-binding proteins that modulate polyadenylation-induced mRNA translation, which is essential for the persistence of memory [19]. CPEBs have been found in several invertebrate and vertebrate genomes, and four *CPEB* genes (*CPEB1–4*) have been identified in mammals [20-24]. All CPEB proteins have two RNA recognition domains (RRM motifs) and a ZZ-type zinc finger domain in the C-terminal region, but differ in their N-terminal domains [25-27]. *Aplysia* CPEB (ApCPEB), *Drosophil*a Orb2, and mouse CPEB3 have two distinct functional conformations that correspond to soluble monomers and amyloidogenic oligomers, and have been implicated in maintenance of long-term facilitation (LTF) in *Aplysia* and long-term memory in both *Drosophil*a and mice [28-34]. In *Drosophil*a, inhibition of amyloid-like oligomerization of Orb2 impairs the persistence of long-lasting memory, and deletion of prion-like domain of Orb2 disrupts long-term courtship memory [32, 35]. The aggregated form of CPEB3, which is inhibited by SUMOylation, can mediate target mRNA translation at activated synapses [36].

Following synaptic stimulation CPEB3 interacts with actin cytoskeleton, with a positive feedback loop of CPEB3/actin regulating remodeling of synaptic structure and connections [37, 38]. Studies of CPEB3 in memory formation revealed that the local protein synthesis and long-term memory storage are regulated by the prion-like CPEB3 aggregates (the aggregation of CPEB3 is thought to strengthen synaptic plasticity in the hippocampus); *CPEB3* conditional knockout mice display impairments in memory consolidation, object placement recognition, and long-term memory maintenance [31]. On the other hand, global *CPEB3* knockout mice display enhanced spatial memory consolidation in the Morris water maze and exhibit elevated short-term fear during the acquisition and extinction of contextual fear memory [39].

The CPEB3 protein is thus well established as a modulator of memory formation and learning, but the function of the CPEB3 ribozyme has not been tested. Given that the self-scission of intronic ribozymes is inversely correlated with splicing efficiency of the harboring pre-mRNA [40], we hypothesized that inhibition of the CPEB3 ribozyme co-transcriptional self-self-cleavage can alter *CPEB3* mRNA splicing and increase the expression of full-length mRNA and CPEB3 protein, leading to polyadenylation of its target mRNAs and an enhancement in the consolidation of hippocampal-dependent memory.

## Results

### CPEB3 mRNA expression and ribozyme activity are upregulated in response to neuronal stimulation

To test this hypothesis, we began by measuring the co-transcriptional self-scission of the murine variant of the ribozyme *in vitro* and found a half-life (t_1/2_) of ∼2 minutes (Fig 1B and Table S1), which is similar to previously measured chimp and fast-reacting human variants of the ribozyme [41]. Because the distance from the ribozyme to the 3^rd^ exon in the *CPEB3* gene is about 10 kb and the RNA polymerase II (RNAPII) transcription rate of long mammalian genes is estimated to be ∼3.5–4.1 knt/min [42], RNAPII would take about 2.5–3 minutes to reach the 3^rd^ exon and mark it for splicing. This result suggested that the ribozyme activity is tuned to the co-transcriptional processing of the CPEB3 pre-mRNA.

The neuronal activity-dependent gene regulation is essential for synaptic plasticity [43]. To investigate the effect of the CPEB3 ribozyme on *CPEB3* mRNA expression and measure its effect on maturation and protein levels, we stimulated primary cortical neurons by glutamate or potassium chloride (KCl). First, *CPEB3* mRNA levels were measured using primers that specifically amplified exon–exon splice junctions (Exons 2–3, 3–6, 6–9; Fig 1A). We found that membrane depolarization by KCl led to an up-regulation of *CPEB3* mRNA 2 hours post stimulation, compared with non-stimulated cultures (Fig 1C). To examine CPEB3 ribozyme activity, total ribozyme and uncleaved ribozyme levels were measured by qRT-PCR, which showed that ribozyme expression is elevated at 1 hour following KCl treatment (Fig 1D). Similarly, glutamate stimulation both increased the expression of spliced exons by 2–3 fold at 2 hours, with a decrease observed at later time points (Fig 1E), and increased ribozyme expression correlated with CPEB3 mRNA expression (Fig 1F). This finding is supported by previous studies showing that synaptic stimulation by glutamate leads to an increase in CPEB3 protein expression in hippocampal neurons [31] and that treatment with kainate likewise induces CPEB3 expression in the hippocampus [21]. The cleaved fraction of the ribozyme was greatest at the highest point of CPEB3 mRNA expression, suggesting efficient co-transcriptional self-scission. Together, these data (i) indicate that the self-cleaving CPEB3 ribozyme is expressed, and potentially activated, in response to neuronal activity, and (ii) suggest that CPEB3 ribozyme *cis*-regulates the maturation of CPEB3 mRNA.

### CPEB3 mRNA and protein levels increase in primary neuronal cultures treated with ribozyme inhibitor

Because our data showed that CPEB3 ribozyme expression is correlated with mRNA expression, we hypothesized that regulation of the ribozyme may modulate CPEB3 mRNA splicing. To test this hypothesis, we inhibited the ribozyme using antisense oligonucleotides (ASOs) spanning the cleavage site (Figs 2A and 2B); these ASOs were similar to those previously used to inhibit *in vitro* co-transcriptional self-scission of this family of ribozymes [13, 44]. ASOs are synthetic single-stranded nucleic acids that can bind to pre-mRNA or mature RNA through base-pairing, and typically trigger RNA degradation by RNase H, thereby turning off the target gene expression. ASOs have also been used to modulate alternative splicing, suggesting that they act co-transcriptionally (e.g., to correct the *SMN2* mRNA) [45]. The ASOs used in this study were designed to increase thermal stability of complementary hybridization and, as a result, to induce higher binding affinity for the ribozyme.

**Fig 2.**
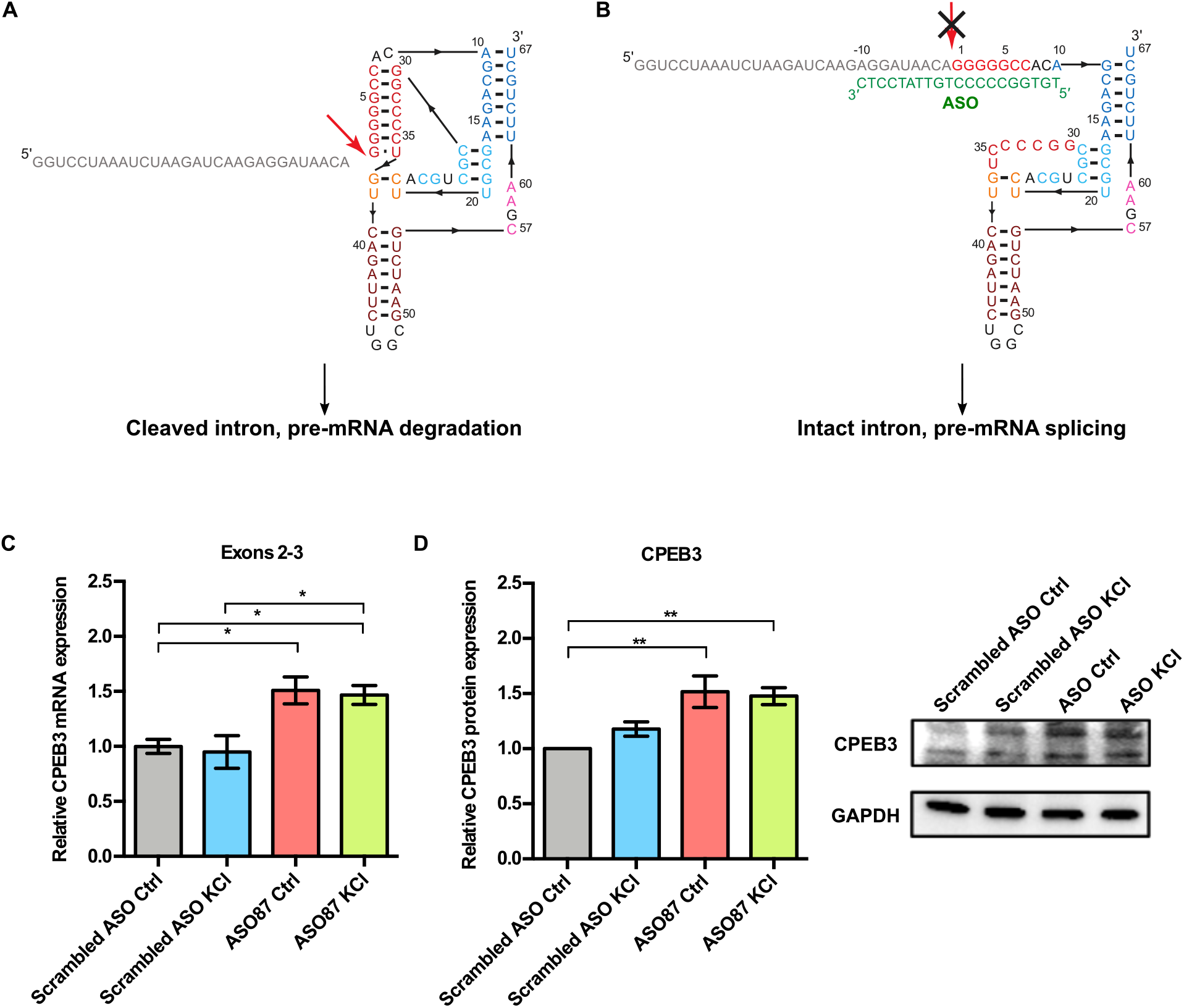
CPEB3 mRNA and protein are upregulated in primary neuronal cultures treated with ASO. (A) Inhibition of the CPEB3 ribozyme by an ASO targeting its cleavage site. Secondary structure of the ribozyme (colored by structural elements [10]). Sequence upstream of the ribozyme is shown in gray, and the site of self-scission is shown with the red arrow. (B) Model of the ribozyme inhibited by the antisense oligonucleotide (ASO, green letters) showing base-pairing between the ASO and 10 nts upstream and downstream of the ribozyme cleavage site. Inhibition of self-scission is indicated by crossed arrow. (C) Ribozyme inhibition by ASO in cultured cortical neurons resulted in upregulation of CPEB3 mRNA (exons 2–3; one-way ANOVA with Sidak’s *post hoc* tests, \**P* < 0.05). (D) Effect of CPEB3 ribozyme ASO on CPEB3 protein expression. GAPDH is used as a loading control (one-way ANOVA with Sidak’s *post hoc* tests, \**P* < 0.05, \*\**P* < 0.01). Data are presented as mean ± SEM.

To study the effect of the CPEB3 ribozyme on *CPEB3* mRNA expression, neuronal cultures were pretreated with either an ASO or a non-targeting control oligonucleotide, followed by KCl stimulation. In the absence of ASO, KCl induced a rapid and robust increase in ribozyme levels compared to cultures containing scrambled ASO, and this effect was suppressed in the presence of ASO, suggesting that the ribozyme is blocked by the ASO (S1A Fig). At an early time point (2 hours post-KCl induction), the ASO-containing culture displayed an increase of spliced mRNA (Figs 2C, S1 B and C), suggesting that the ASO prevents CPEB3 ribozyme from cleaving the intron co-transcriptionally and promotes mRNA maturation. At 24 hours post-KCl induction, we observed no significant difference in CPEB3 ribozyme expression among groups (S1D Fig). Likewise, the level of *CPEB3* mRNA exons 2–3 returned to the basal level (S1E Fig), while exons 3–6 remained slightly elevated in the ASO-treatment groups (S1F Fig). The mRNA expression of CPEB3 exons 6–9 remained stable over time and was not affected by ASO treatment or KCl stimulation (S1G Fig). Taken together, these data suggest that the CPEB3 ribozyme modulates the production of the full-length *CPEB3* mRNA.

To evaluate whether the ASO specifically targets CPEB3 ribozyme or modulates intron levels in general, we measured the levels of the 4^th^ CPEB3 intron, which does not harbor a self-cleaving ribozyme. No significant difference in the 4^th^ intron expression between groups was observed, demonstrating that the ASO does not have a broad non-specific effect on the stability of other introns (Fig S2H). Furthermore, to assess whether the ASO induces cytotoxicity *in vitro*, neuronal cultures were treated with either ASO or scrambled ASO. Cell viability was measured with an XTT assay, revealing no difference in either ASO- or scrambled ASO-treated cells, compared to untreated cells. These data suggest that the ASOs used in this study did not induce cytotoxic effects in cultured neurons (S1I Fig).

We next determined whether inhibition of CPEB3 ribozyme regulates CPEB3 protein expression. Treatment with the ribozyme ASO resulted in a significant increase in CPEB3 protein levels in both the basal state and under KCl-stimulated conditions, indicating a coordination of activity-dependent transcription and translation upon inhibition of CPEB3 ribozyme activity (Fig 2D).

### Ribozyme inhibition leads to increased expression of plasticity-related proteins

In *Aplysia* sensory-motor neuron co-culture, application of repeated pulses of serotonin (5-HT) induces ApCPEB protein expression at the stimulated synapses and LTF, which is a form of learning-related synaptic plasticity that is widely studied in *Aplysia* [20, 29]. In murine primary hippocampal neurons, the level of CPEB3 protein expression is positively regulated by neuronal activity [31] and plays dual roles in regulating mRNA translation [37, 46] whereby a post-translational modification of CPEB3 can convert it from a repressor to an activator: a monoubiquitination by Neuralized1 leads to activation of CPEB3, which promotes subsequent polyadenylation and translation of GluA1 and GluA2 [47]. Previous studies have also demonstrated the role of CPEB3 in the translational regulation of a number of plasticity-related proteins (PRPs), including AMPA-type glutamate receptors (AMPARs), NMDA receptor (NMDAR), and postsynaptic density protein 95 (PSD-95) [26, 31, 39, 48]. As an RNA binding protein, CPEB3 has been shown to bind to 3*’* UTR of GluA1, GluA2, and *PSD-95* mRNAs and to regulate their polyadenylation and translation [26, 31, 39, 47].

To test whether inhibition of CPEB3 ribozyme modulates expression of PRPs, we measured the protein levels. We found that under KCl-induced depolarizing conditions, treatment with the CPEB3 ribozyme ASO resulted in a significant increase in GluA1 and PSD-95 protein expression, whereas GluA2 levels remained unchanged (S2A and S2B Fig). Likewise, ASO treatment led to an up-regulation of NR2B protein, one of the NMDAR subunits (S2C and S2D Fig). These results suggest that CPEB3 ribozyme activity affects several downstream processes, particularly mRNA maturation and translation, as well as the expression of PRPs, including the translation of AMPAR and NMDAR mRNAs.

### CPEB3 ribozyme ASO leads to an increase of CPEB3 mRNA and polyadenylation of PRPs in the CA1 hippocampus

To investigate whether the CPEB3 ribozyme exhibits similar effects in regulating genes related to synaptic plasticity *in vivo*, mice were stereotaxically infused with either ribozyme ASO, scrambled ASO, or vehicle into the CA1 region of the dorsal hippocampus, a major brain region involved in memory consolidation and persistence (Fig 3A). Infusion of ASO targeting the CPEB3 ribozyme significantly reduced ribozyme levels detected by RT-qPCR in the dorsal hippocampus (S3A Fig). We found that administration of ASO led to an increase of *CPEB3* mRNA in the CA1 hippocampus (Fig 3B), confirming that the ASO prevents ribozyme self-scission during CPEB3 pre-mRNA transcription, thereby increasing *CPEB3* mRNA levels. No significant difference in the ribozyme-free 4^th^ intron levels was observed between ASO and vehicle (S3B Fig).

**Fig 3.**
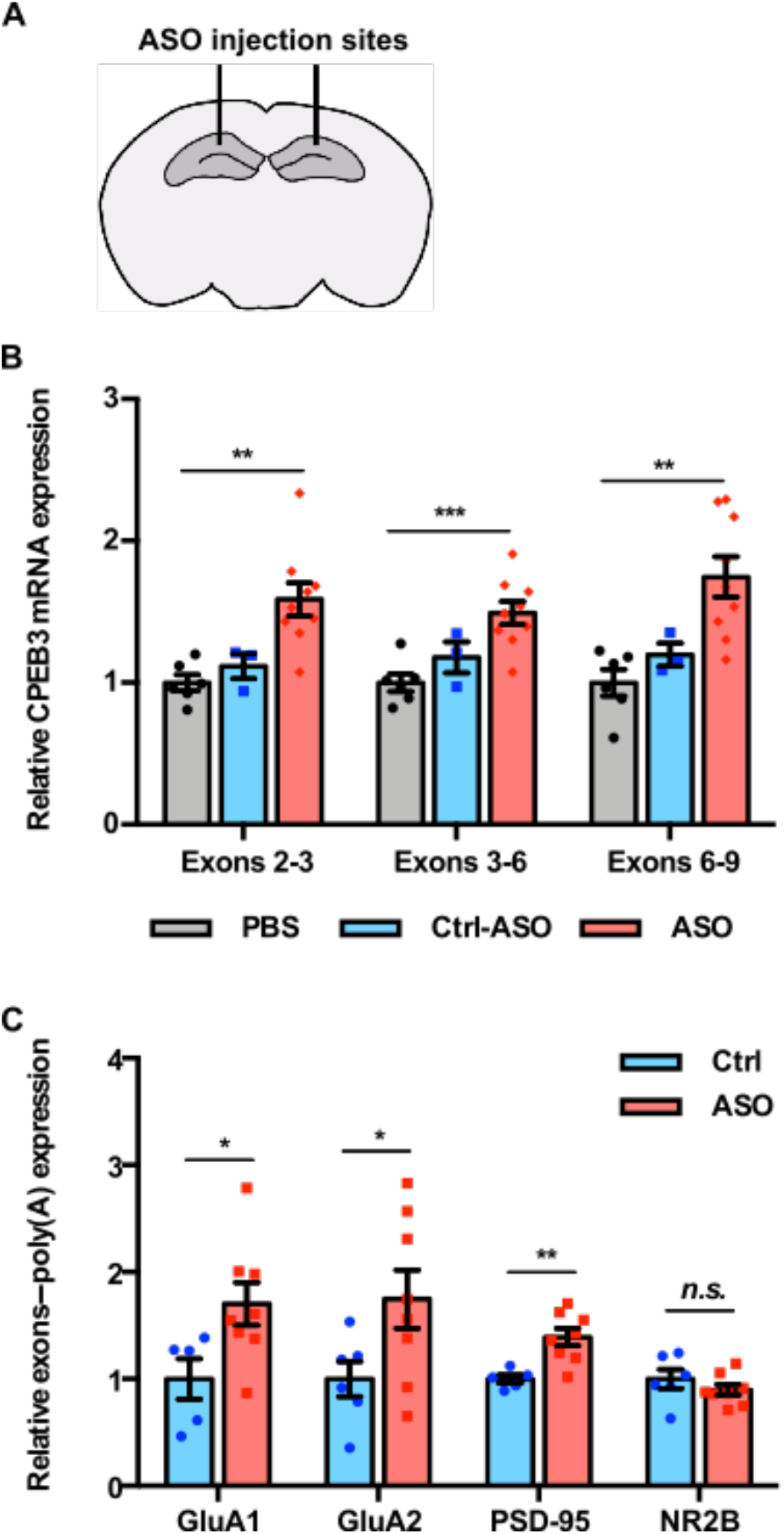
CPEB3 ribozyme ASO leads to an increase of CPEB3 mRNA and polyadenylation of PRPs in the CA1 hippocampus. (A) Schematic representation of stereotaxic procedure. ASO, scrambled ASO, or vehicle was bilaterally infused to the mouse CA1 hippocampus. (B) CPEB3 mRNA expression is upregulated in the CPEB3 ribozyme ASO treatment group compared to controls (one-way ANOVA with Sidak’s *post hoc* tests. \*\**P* < 0.01, \*\*\**P* < 0.001). (C) Inhibition of CPEB3 ribozyme results in increased polyadenylation of plasticity-related genes (unpaired *t* test, \**P* < 0.05, \*\**P* < 0.01, *n*.*s*. not significant). Data are presented as mean ± SEM.

Next, we tested whether the CPEB3 ribozyme inhibition affects CPEB3 translation, and no significant difference between ASO and control groups was observed (S5A and S5B Fig). We further observed that blocking the CPEB3 ribozyme does not change GluA1, GluA2, PSD-95, and NR2B mRNA or protein expression in naïve mice (S4, S5A and S5C Fig). Thus, in naïve mice, ribozyme inhibition leads to increased basal levels of the *CPEB3* mRNA, but the levels of the CPEB3 protein and its downstream mRNA targets remain unchanged.

To further delineate whether the CPEB3 ribozyme activity results in polyadenylation of its target mRNAs, 3′ rapid amplification of cDNA ends (3′ RACE) was performed to examine the 3′ termini of several mRNAs. We found that ribozyme ASO administration led to increased GluA1, GluA2, and PSD-95 mRNA polyadenylation in the mouse dorsal hippocampus (Fig 3C). These data support a model wherein the inhibition of the CPEB3 ribozyme leads to increased polyadenylation of existing AMPARs and PSD-95 mRNAs, and suggests a role in post-transcriptional regulation and 3′ mRNA processing.

### Inhibition of CPEB3 ribozyme in the dorsal hippocampus enhances long-term memory

To assess whether inhibition of the CPEB3 ribozyme improves memory formation, we tested it with respect to long-term memory using the object location memory (OLM) task (Fig 4A). The OLM task has been widely used to study hippocampal-dependent spatial memory. The task is based on an animal’s innate preference for novelty and its capability for discriminating spatial relationships between novel and familiar locations [49]. During a testing session, mice retrieve the memory that encoded for the objects they were exposed to in the training session. We infused mice bilaterally into the CA1 dorsal hippocampus with the CPEB3 ribozyme ASO, scrambled ASO, or vehicle 48 hours prior to OLM training. The CPEB3 ribozyme ASO group showed a significant increase in discrimination index (DI) between training and testing compared to control groups, suggesting that these mice experienced a robust enhancement of novel object exploration (Fig 4B). We observed no significant difference in training DI (*P* > 0.05), indicating that mice exhibit no preference for either object (Fig 4B). Likewise, during training and testing sessions, ASO-infused mice and control mice displayed similar total exploration time, demonstrating that both groups of mice have similar exploitative behavior (Fig 4C). These results provide strong evidence that CPEB3 is critical for long-term memory, and that the CPEB3 ribozyme activity is anti-correlated with the formation of long-term memory.

**Fig 4.**
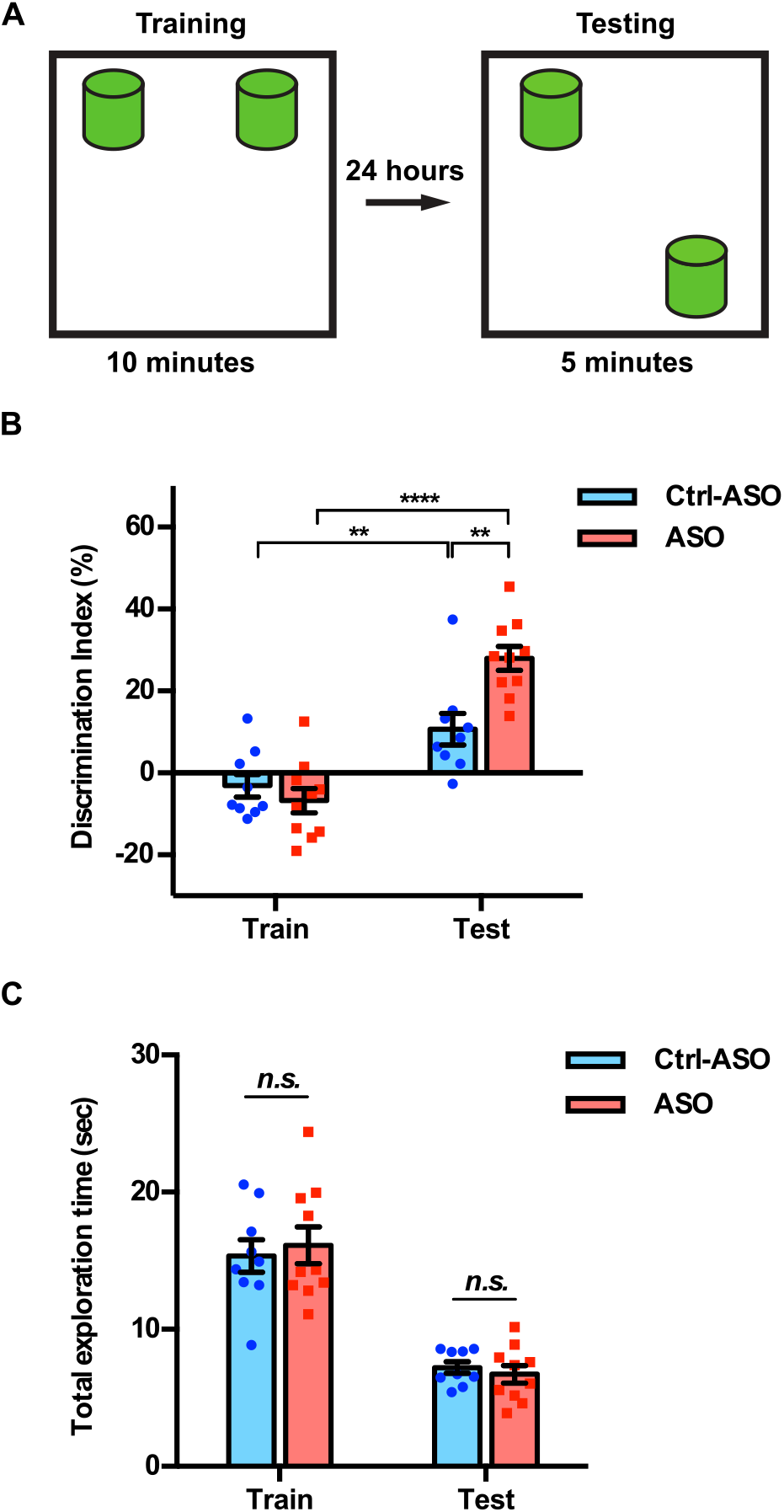
Inhibition of CPEB3 ribozyme enhances long-term OLM. (A) Experimental procedure testing long-term memory. (B) Mice infused with CPEB3 ribozyme ASO show significant discrimination index in OLM testing (two-way ANOVA with Sidak’s *post hoc* tests, \**P* < 0.05, \*\**P* < 0.01, \*\*\*\**P* < 0.0001). (C) CPEB3 ribozyme ASO and control mice display similar total exploration time (one-way ANOVA with Sidak’s *post hoc* tests, *n*.*s*. not significant). Data are presented as mean ± SEM.

### CPEB3 ribozyme ASO leads to an upregulation in protein expression of CPEB3 and PRPs during memory consolidation

Learning-induced changes in gene expression and protein synthesis are essential for memory formation and consolidation [50]. To determine whether upregulation of CPEB3 mRNA by the ribozyme ASO leads to a change in expression of the CPEB3 protein and its downstream targets, we analyzed the dorsal hippocampal homogenates and synaptosomal fractions. Administration of CPEB3 ribozyme ASO led to a significant increase of CPEB3 protein expression in the CA1 hippocampal homogenates and crude synaptosomes 1 hour after OLM testing (Figs 5, A, B, and D). This result confirms that ASO-mediated knockdown of the CPEB3 ribozyme facilitates CPEB3 mRNA processing and translation. The protein levels of GluA1, GluA2, PSD-95, and NR2B were measured to determine whether increased CPEB3 further regulates translation of PRPs. In total tissue lysates, no significant difference in PRPs levels was observed between ASO and control (Figs 5, A and C). However, in synaptosomal fractions, GluA1, PSD-95, and NR2B protein levels were increased in ASO-infused mice, relative to scrambled ASO control animals; and GluA2 protein level remained unaffected (Figs 5, A and E). Our findings thus show that blocking CPEB3 ribozyme activity leads to an increase in CPEB3 protein production, and up-regulation of CPEB3 by OLM further mediates local GluA1, PSD-95, and NR2B translation.

**Fig 5.**
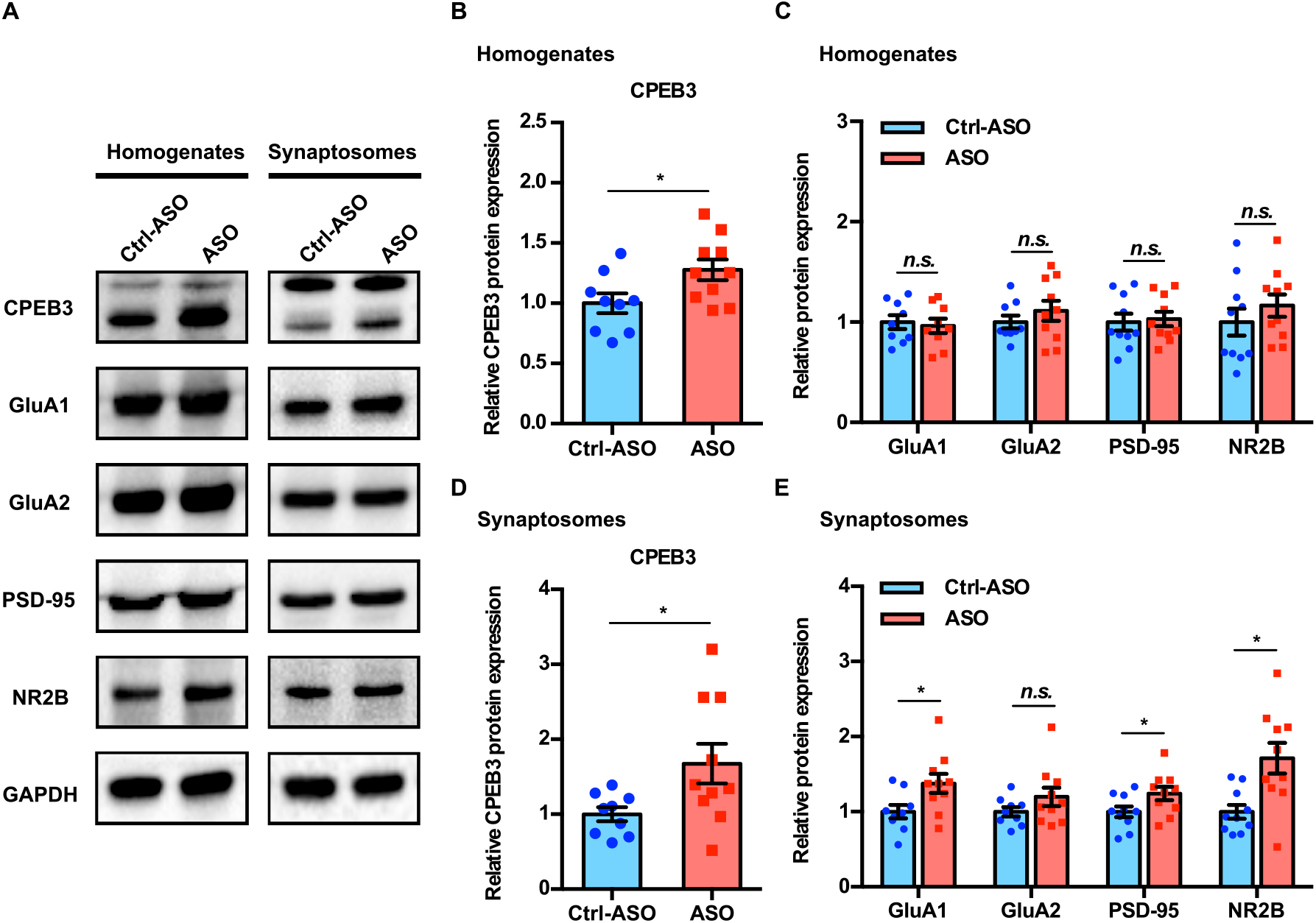
Inhibition of CPEB3 ribozyme leads to upregulation of CPEB3 and PRPs protein expression after OLM. (A) Representative images of immunoblotting analysis. GAPDH is used as a loading control. Quantification of CPEB3 (B) and PRPs (C) in tissue homogenates shows increased CPEB3, but not PRPs, protein expression (unpaired *t* test, \**P* < 0.05, *n*.*s*. not significant). (D) In synaptosomes, the protein expression of both CPEB3 (D) and PRPs (E) is increased (unpaired *t* test, \**P* < 0.05, *n*.*s*. not significant). Data are presented as mean ± SEM.

## Discussion

Self-cleaving ribozymes are broadly distributed small functional RNAs that catalyze a site-specific scission of their backbone [1]. The HDV family of these ribozymes acts during rolling circle replication of the HDV RNA genome and in processing of retrotransposons [2, 3, 14-16], but given their broad distribution in nature, their biological roles are largely unexplored. Mammals harbor several self-cleaving ribozymes—all without known biological functions [4-9]. One of these ribozymes, the HDV-like CPEB3 ribozyme, maps to the second intron of the *CPEB3* gene and its *in vitro* activity (Figs 1A and 1B) suggested that its rate of self-scission may be tuned to disrupt the intron at a rate that is similar to the production speed of the downstream intronic sequence ahead of the next exon. Previous work on synthetic ribozymes placed within introns of mammalian genes showed that splicing of the surrounding exons is sensitive to the continuity of the intron: fast ribozymes caused efficient self-scission of the intron, leading to unspliced mRNA and resulting in lower protein expression, whereas slow ribozymes had no effect on mRNA splicing and subsequent protein expression [40]. Inspired by this work, we investigated how the intronic ribozyme affects the *CPEB3* mRNA maturation and translation, and its effect on memory formation in mice.

Modifications of synaptic strength are thought to underlie learning and memory in the brain. Studies in hippocampal slices revealed local translation in dendrites following induction of LTP [51]. Cytoplasmic polyadenylation-induced translation is one of the key steps for regulating protein synthesis and neuroplasticity [22, 46, 52]. One of the proteins involved in regulation of cytoplasmic polyadenylation of mRNAs is CPEB3. Recent studies have shown that CPEB3 regulates mRNA translation of several PRPs at synapses, where it is essential for synaptic strength [26, 31, 47]. Previous reports have shown that CPEB3 regulates GluA1 and GluA2 polyadenylation: CPEB3 conditional knockout mice fail to elongate the poly(A) tail of GluA1 and GluA2 mRNA after Morris water maze training, and overexpression of CPEB3 changes the length of the GluA1 and GluA2 mRNA poly(A) tail [31]. Even though translational control by regulation of CPEB3 has been demonstrated to contribute to the hippocampal-dependent learning and memory [47], one unaddressed question is whether the CPEB3 expression is modulated by the CPEB3 ribozyme. In mammals, the coordination of pre-mRNA processing and transcription can affect its gene expression [53]. Recent measurement of co-transcriptional splicing events in mammalian cells using long-read sequencing and Precision Run-On sequencing (PRO-seq) approaches demonstrated that co-transcriptional splicing efficiency impacts productive gene output [54]. The temporal and spatial window shows that the splicing and transcription machinery are tightly coupled. In agreement with this co-transcriptional splicing model, our study shows that inhibition of the intronic CPEB3 ribozyme leads to an increase in *CPEB3* mRNA and protein levels in primary cortical neurons and the dorsal hippocampus upon synaptic stimulation, and leading to changes in polyadenylation of target mRNAs of the CPEB3 protein.

Activity-dependent synaptic changes are governed by AMPAR trafficking, and AMPARs are mobilized to the post-synaptic surface membrane in response to neuronal activity in a dynamic process [55]. Our data demonstrate that the activation of CPEB3 by neuronal stimulation further facilitates translation of PRPs *in vivo*. These observations are consistent with a model in which learning induces CPEB3 protein expression, and ablation of CPEB3 abolishes the activity-dependent translation of GluA1 and GluA2 in the mouse hippocampus [31]. Specifically, it has been suggested that CPEB3 converts to prion-like aggregates in stimulated synapses that mediate hippocampal synaptic plasticity and facilitate memory storage [56]. Because training can produce effective long-term memory, it is likely that increased CPEB3 protein expression due CPEB3 ribozyme inhibition further facilitates experience-induced local translational processes.

ASOs have been used in many studies to inhibit specific mRNAs. A notable example is an FDA-approved ASO that modulates co-transcriptional splicing of the *SMN2* mRNA [45]. Our works shows that an ASO designed to bind the substrate strand of an endogenous self-cleaving ribozyme located in an intron increases the expression of the fully spliced mRNA that harbors the ribozyme. Interestingly, experiments with inhibitory ASO yielded lower the ribozyme levels than control experiments, suggesting that the ASO directs degradation of the target sequence; however, this degradation must occur on a timescale that is longer than the splicing of the mRNA, because we consistently see higher mRNA levels when the ribozyme is inhibited. Further studies will be necessary to delineate the full mechanism of action of anti-ribozyme ASOs. Given that three endogenous mammalian self-cleaving ribozymes map to introns [4, 6, 7], we anticipate that application of ASOs will help decipher their effect on their harboring mRNAs and establish their biological roles.

In summary, we have delineated an important step in molecular mechanisms underlying a unique role for the CPEB3 ribozyme in post-transcriptional maturation of CPEB3 mRNA and its subsequent translation in mouse CA1 hippocampus. Inhibition of the CPEB3 ribozyme by ASO and OLM training induce activity-dependent upregulation of CPEB3 and local production of PRPs. These molecular changes are critical for establishing persistent changes in synaptic plasticity that are required for long-term memory, and represent a biological role for self-cleaving ribozymes in the brain. More broadly, our study demonstrates a novel biological role for self-cleaving ribozymes and the first example of their function in mammals.

## Materials and Methods

### Primary cortical neuronal culture

Pregnant female C57BL/6 mice (The Jackson Laboratory) were euthanized at E18, and embryos were collected into an ice-cold Neurobasal medium (Thermo Fisher Scientific). Embryonic cortices were dissected, meninges were removed, and tissues were minced. Cells were mechanically dissociated, passed through a 40-µm cell strainer, counted, and plated at a density of 0.5 × 10^6^ cells per well in six-well plates coated with poly-D-lysine (Sigma-Aldrich). Neuronal cultures were maintained at 37 °C with 5% CO_2_, and grown in Neurobasal medium containing 2% B27 supplement (Thermo Fisher Scientific), 1% penicillin/streptomycin (Thermo Fisher Scientific), and 2 mM L-Glutamine (Thermo Fisher Scientific) for 7–10 days *in vitro* (DIV), with 50% of the medium being replaced every 3 days. All experimental procedures were performed according to the National Institutes of Health Guide for the Care and Use of Laboratory Animals and approved by the Institutional Animal Care and Use Committee of the University of California, Irvine.

### Mice

C57BL/6J mice (8–10 weeks old, The Jackson Laboratory) were housed in a 12-h light/dark cycle and had free access to water and food. All experiments were conducted during the light cycle. All experimental procedures were performed according to the National Institutes of Health Guide for the Care and Use of Laboratory Animals and approved by the Institutional Animal Care and Use Committee of the University of California, Irvine.

### Measurement of co-transcriptional self-scission of the CPEB3 ribozyme

Transcription reactions were set up in a 5-µL volume and incubated for 10 minutes at 25 °C, as described previously [17]. The reactions contained: 1 µL of 5x transcription buffer (10 mM spermidine, 50 mM dithiothreitol, 120 mM Tris chloride buffer, pH 7.5, and 0.05% Triton X-100), 1 µL of 5x ribonucleoside triphosphates (final concentration of 6.8 mM), 1 µL of 5 mM Mg^2+^, 1 µL DNA amplified by PCR to about 1 µM final concentration, 0.5 µL of 100% DMSO, 0.15 µL of water, 0.1 µL of murine RNase inhibitor (40,000 units/mL, New England Biolabs), 0.125 µL of T7 polymerase, and 0.125 µL [α-^32^P]ATP. To prevent initiation of new transcription, the reactions were diluted into 100 µL of physiological-like buffer solution at 37 °C. The solution consisted of 2 mM Mg^2+^ (to promote ribozyme self-scission), 140 mM KCl, 10 mM NaCl, and 50 mM Tris chloride buffer (pH 7.5). The 100-µL solution was then held at 37 °C for the reminder of the experiment while aliquots were withdrawn at various time points. An equal volume of 4 mM EDTA/7 M urea stopping solution was added to each aliquot collected. Aliquots were resolved using denaturing polyacrylamide gel electrophoresis (PAGE, 7.5% polyacrylamide, 7 M urea) at 20 W. The PAGE gel was exposed to a phosphosimage screen for ∼2 hours and analyzed using a Amersham Typhoon imaging system (GE Healthcare). Band intensities corresponding to the uncleaved ribozymes and the two products of self-scission were analyzed using ImageQuant (GE Healthcare) and exported into Excel. Fraction Intact was calculated as the intensity of the band corresponding to the uncleaved ribozyme divided by the sum of band intensities in a given PAGE lane. The data were fit to a biexponential decay model:

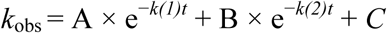

In the case of the minimum murine CPEB3 ribozyme construct (−10/72; S2 Fig and Table S1), the data were modeled by a monoexponential decay with an uncleaved fraction (using parameters A, *k*_1_, and C only).

### Antisense oligonucleotides (ASOs)

ASOs used in this study are 20 nucleotides in length and are chemically modified with 2′-*O*-methoxyethyl (MOE, underlined) and 2′-4′ constrained ethyl (cEt, bold) [57]. All internucleoside linkages are modified with phosphorothioate linkages to improve nuclease resistance. ASOs were solubilized in sterile phosphate-buffered saline (PBS). The sequences of the ASOs are as follows (all cytosine nucleobases are 5-methyl substituted):

Scrambled control ASO: 5′-C<u>CT</u>T<u>CC</u>C<u>TG</u>A<u>AG</u>G<u>TT</u>C<u>CT</u>C<u>C</u>-3′;

CPEB3 ribozyme ASO: 5′-T<u>GT</u>G<u>GC</u>C<u>CC</u>C<u>TG</u>T<u>TA</u>T<u>CC</u>T<u>C</u>-3′.

### Neuronal stimulation

Neurons were treated with ASO or scrambled ASO (1 µM) for 18 hours prior to neuronal stimulation. To study activity-dependent gene regulation, neuronal cultures were treated with vehicle, 5 µM glutamate (10 minutes), or 35 mM KCl (5 minutes). After stimulation, cultures were washed with Hanks’ buffered salt solution (HBSS, Thermo Fisher Scientific), and then replaced with fresh medium.

### Quantitative RT-PCR analysis

Total RNA was isolated from primary cortical neurons or mouse hippocampus using TRI reagent (Sigma-Aldrich) according to the manufacturer’s protocol. RNA concentration was measured using a NanoDrop ND-1000 spectrophotometer (Thermo Fisher Scientific). Total RNA was reverse transcribed using random decamers and M-MLV reverse transcriptase (Promega)/Superscript II RNase H reverse transcriptase (Thermo Fisher Scientific). Quantitative RT-PCR was performed on a BioRad CFX Connect system using iTaq Universal SYBR Green Supermix (BioRad). Designed primers were acquired from Integrated DNA Technologies and provided in Table S2. Desired amplicons were verified by melting curve analysis and followed by gel electrophoresis. The starting quantity of DNA from each sample was determined by interpolation of the threshold cycle (CT) from a standard curve of each primer set. Relative gene expression levels were normalized to the endogenous gene *GAPDH*.

### Immunoblotting

Primary cortical neurons or mouse hippocampal tissues were lysed in RIPA lysis buffer with protease inhibitor (Santa Cruz Biotechnology). Crude synaptosomal fractions were prepared as previously described [58]. Protein concentrations were measured using bicinchoninic acid (BCA) protein assay (Thermo Fisher Scientific). Ten to 30 µg of protein samples were loaded on 10% sodium dodecyl sulfate polyacrylamide (SDS-PAGE) gels and separated by electrophoresis. Gels were electro-transferred onto polyvinylidene fluoride (PVDF) membranes using a semi-dry transfer system (BioRad). Membranes were either blocked with 5% nonfat milk or 5% bovine serum albumin (BSA) in Tris-buffered saline/Tween 20 (0.1% [vol/vol]) (TBST) for 1 hour at room temperature. Membranes were incubated with primary antibodies overnight at 4 °C. After primary antibody incubation, membranes were washed three times with TBST and then incubated with secondary antibodies for 1 hour at room temperature. Bands were detected using an enhanced chemiluminescence (ECL) kit (Thermo Fisher Scientific), visualized using BioRad Chemidoc MP imaging system, and analyzed by Image Lab software (BioRad). GAPDH was used as a loading control. All antibodies used in this study are listed in Table S3.

### *In vitro* XTT cell viability assay

Primary cortical neurons (10,000 to 20,000 cells/well) were plated onto 96-well plates coated with poly-D-lysine. After 7–14 days, ASOs or scrambled ASOs were added and incubated for 18 hours. Cell viability was determined using the 2,3-bis[2-methoxy-4-nitro-5-sulfophenyl]-2H-tetrazolium-5-carboxyanilide inner salt (XTT) assay according to the manufacturer’s protocol (Biotium). The assay utilizes the ability of viable cells with active metabolism to reduce the yellow tetrazolium salt to the soluble orange formazan product by mitochondrial dehydrogenase enzymes. The XTT reagent was added to each well and incubated for 2–4 hours at 37 °C and 5% CO_2_. Absorbance was measured at 450 nm with a reference wavelength of 680 nm using a Biotek Synergy HT microplate reader. Results were normalized to control, and all samples were assayed in triplicate.

### Stereotaxic surgeries

C57/BL6J mice (8–10 weeks old, Jackson Laboratory), housed under standard conditions with light-control (12-h light/12-h dark cycles), were anaesthetized with an isoflurane (1–3%)/oxygen vapor mixture. Mice were infused bilaterally to the CA1 region of the dorsal hippocampus with ribozyme ASO, scrambled ASO diluted in sterile PBS, or vehicle. The following coordinates were used, relative to bregma: medial-lateral (ML), ±1.5 mm; anterior-posterior (AP), −2.0 mm; dorsal-ventral (DV), −1.5 mm. ASOs or vehicle (1 nmol/µL) were infused bilaterally at a rate of 0.1 µL/min using a Neuros Hamilton syringe (Hamilton company) with a syringe pump (Harvard Apparatus). The injectors were left in place for 2 minutes to allow diffusion, and then were slowly removed at a rate of 0.1 mm per 15 sec. The incision site was sutured, and mice were allowed to recover on a warming pad and then were returned to cages. For all surgeries, mice were randomly assigned to the different conditions to avoid grouping same treatment conditions in time.

### Object location memory (OLM) tasks

The OLM was performed to assess hippocampus-dependent memory, as previously described [49]. Briefly, naïve C57/BL6J mice (8–12 weeks old; n = 10–12/group; ribozyme ASO, scrambled ASO) were trained and tested. Prior to training, mice were handled 1–2 minutes for 5 days and then habituated to the experimental apparatus for 5 minutes on 6 consecutive days in the absence of objects. During training, mice were placed into the apparatus with two identical objects and allowed to explore the objects for 10 minutes. Twenty-four hours after training, mice were exposed to the same arena, and long-term memory was tested for 5 minutes, with the two identical objects present, one of which was placed in a novel location. For all experiments, objects and locations were counterbalanced across all groups to reduce bias. Videos of training and testing sessions were analyzed for discrimination index (DI) and total exploration time of objects. The videos were scored by observers blind to the treatment. The exploration of the objects was scored when the mouse’s snout was oriented toward the object within a distance of 1 cm or when the nose was touching the object. The relative exploration time was calculated as a discrimination index (DI = (*t*_novel_ – *t*_familiar_) / (*t*_novel_ + *t*_familiar_) × 100%). Mice that demonstrated a location or an object preference during the training trial (DI > ±20) were removed from analysis.

### 3’ RACE

Total RNA was extracted from the mouse CA1 hippocampus. 3*’* rapid amplification of cDNA ends (3*’* RACE) was performed to study the alternative polyadenylation. cDNA was synthesized using oligo(dT) primers with 3*’* RACE adapter primer sequence at the 5*’* ends. This cDNA library results in a universal sequence at the 3*’* end. A gene-specific primer (GSP) and an anchor primer that targets the poly(A) tail region were used for the first PCR using the following protocol: 95 °C for 3 minutes, followed by 30 cycles of 95 °C for 30 seconds, 55 °C for 30 seconds, and 72 °C for 3 minutes, with a final extension of 72 °C for 5 minutes. To improve specificity, a nested PCR was then carried out using primers internal to the first two primers. Upon amplification condition optimization, a quantitative PCR was performed on the first diluted PCR product using the nested primers, and a standard curve of the primer set was generated to determine the effect of relative expression of 3*’*-mRNA and alternative polyadenylation. All primers used in this study are listed in Table S4. When resolved using agarose gel electrophoresis, this nested-primer qPCR produced single bands corresponding to the correct amplicons of individual cDNAs.

### Statistical analysis

Data are presented as means ± SEM. Statistical analyses were performed using GraphPad Prism (GraphPad Prism Software). Statistical differences were determined using two-tailed Welch’s *t* test when comparing between 2 independent groups, and one-way ANOVA with Sidak’s *post hoc* tests when comparing across 3 or more independent groups. OLM data were analyzed with two-way ANOVA followed by Sidak’s *post hoc* tests. *P* < 0.05 was considered significant.

## Acknowledgments

We thank M. Malgowska, C-K. Lau, and M. A. Sta Maria for experimental assistance.

## Funding

National Institutes of Health grant R01AG051807 (MAW)

National Institutes of Health grant RF1AG057558 (MAW)

National Science Foundation 1804220 (AL)

National Science Foundation 1330606 (AL)

National Science Foundation Graduate Research Fellowship (CCC)

## Author contributions

Design of cell culture experiments: CCC, XL, TWB, AL

Design of mouse experiments: CCC, MAW, AL

In vitro ribozyme kinetics measurements: MM

Design of ASOs: MN

Cell culture experiments: CCC, LT, XL

Mouse experiments: CCC

Stereotaxic surgeries and in vivo behavior experiments: JH, CC

Data analysis: CCC

Writing—original draft: CCC, AL

Writing—review & editing: CCC, MN, XL, LT, TWB, MAW, AL

## Competing interests

All other authors declare they have no competing interests.

## Data and materials availability

All data are available in the main text or the supplementary materials.

## Supplementary Materials for

**S1 Fig.**
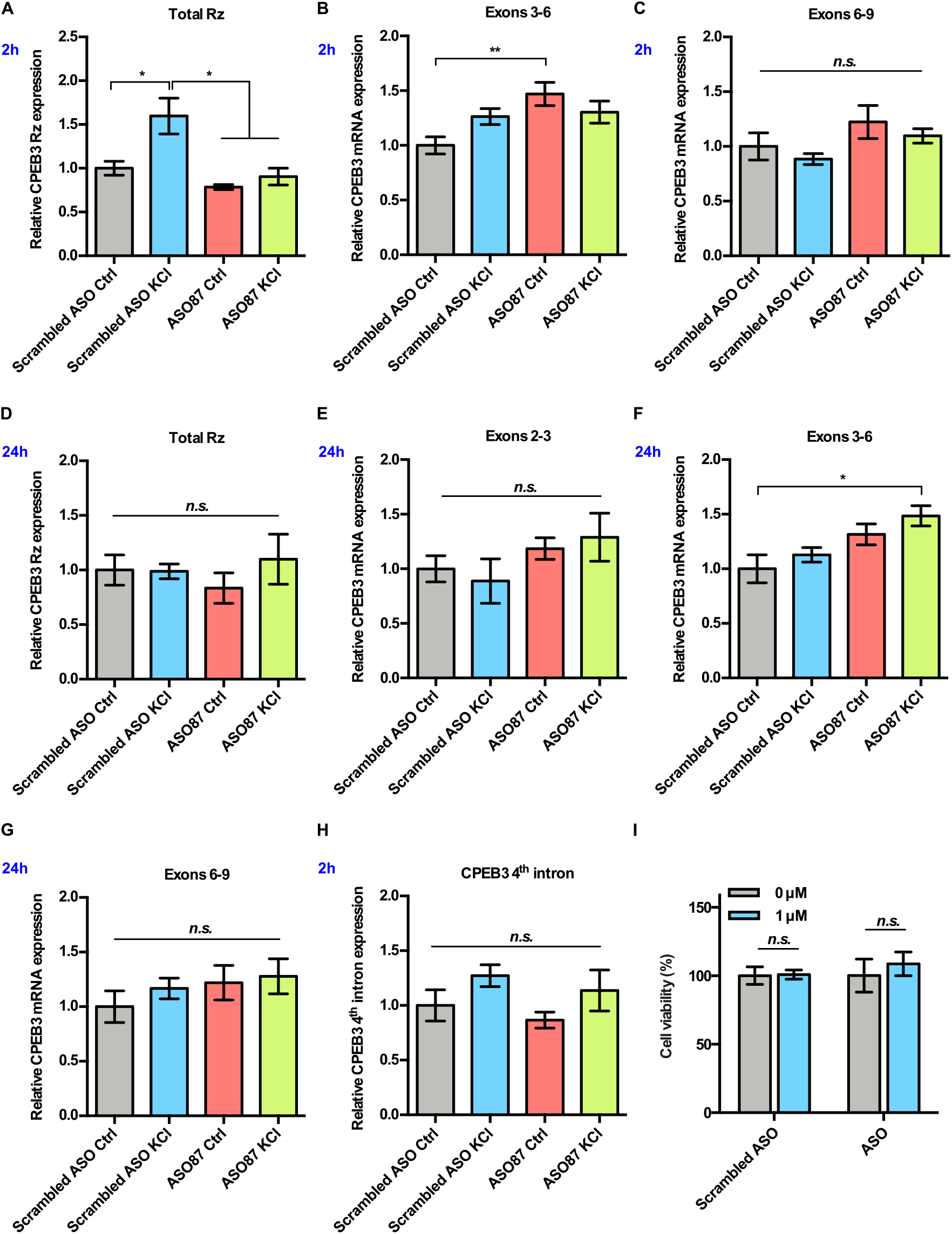
Effect of CPEB3 ribozyme ASO on CPEB3 expression in embryonic cortical neurons. (A) CPEB3 ribozyme levels increase together with levels of the surrounding exons 2 hours post stimulation in experiments with control ASO. Ribozyme levels are significantly lower in ribozyme ASO experiments, suggesting blocking of the RT-PCR reaction by the ASO. (B) and (C) Inhibition of CPEB3 ribozyme by ASO resulted in upregulation of CPEB3 mRNA basal levels for exons 3–6 (B) at 2-hour time point. Levels of exons 6–9 did not increase significantly (C) (one-way ANOVA with Sidak’s *post hoc* tests, \*\**P* < 0.01, *n*.*s*. not significant). (D) No statistically significant difference in CPEB3 ribozyme expression was observed after 24 hours post KCl induction (one-way ANOVA with Sidak’s *post hoc* tests, *n*.*s*. not significant), suggesting that all intronic RNA levels reached basal levels. (E) – (G) CPEB3 mRNA expression largely returned to the basal level 24 hours post stimulation, although levels of spliced exons 3–6 remained elevated (E: exons 2–3, F: exons 3–6, G: exons 6–9, one-way ANOVA with Sidak’s *post hoc* tests, \**P* < 0.05, *n*.*s*. not significant). (H) qRT-PCR analysis of CPEB3 4^th^ intron expression reveals that the ribozyme ASO does not affect its levels, suggesting that it is specific for the ribozyme (one-way ANOVA with Sidak’s *post hoc* tests, *n*.*s*. not significant). (I) Effect of ASOs treatment on cell viability. XTT assay was performed after 18 hours incubation of ASOs. Relative cell viability was normalized to the vehicle control (unpaired *t* test, *n*.*s*. not significant). Data are presented as mean ± SEM.

**S2 Fig.**
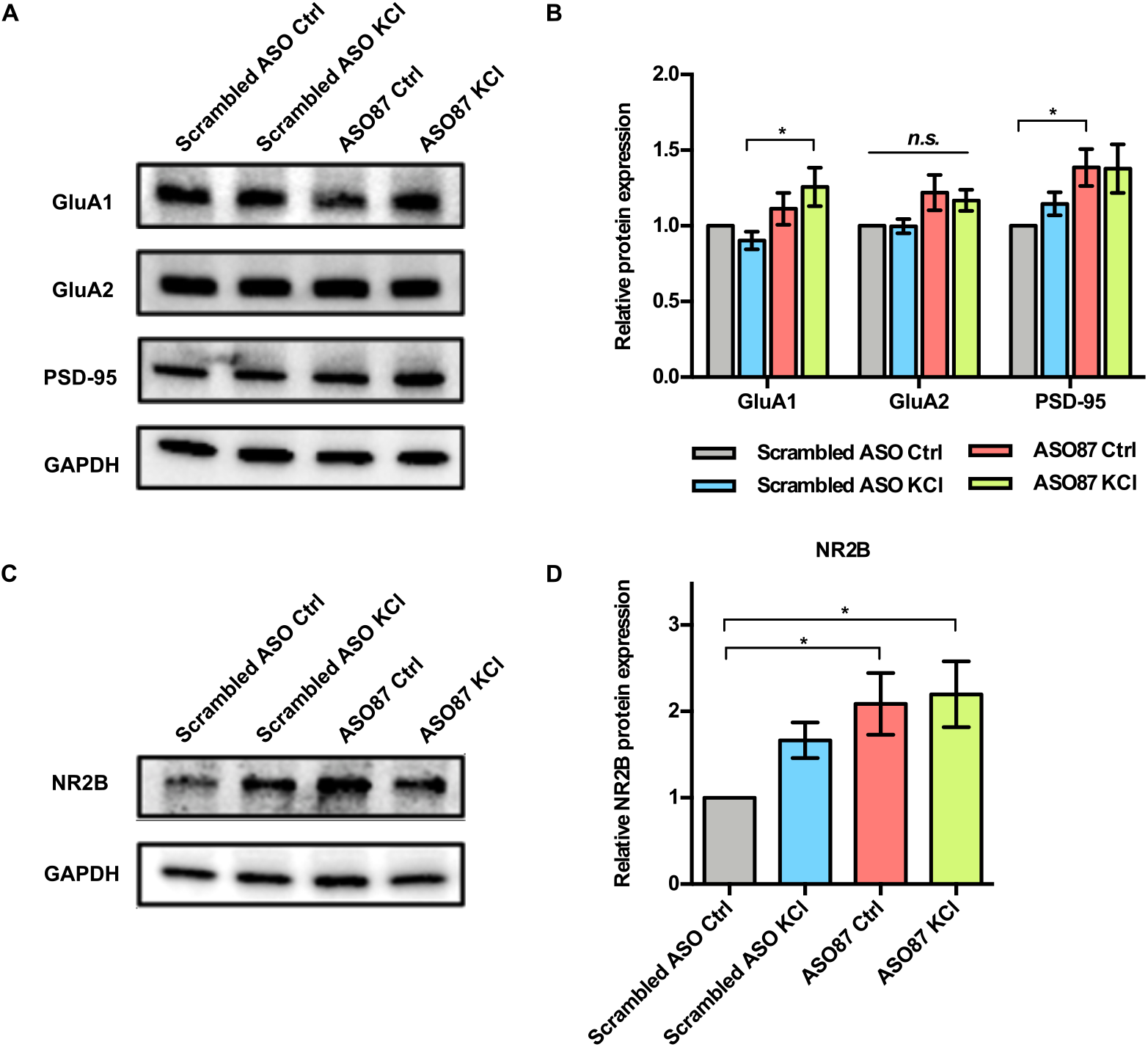
Effect of CPEB3 ribozyme ASO on protein expression in embryonic cortical neurons. Primary neuronal cultures were pretreated with ASO or scrambled ASO, followed by KCl stimulation. Cells were harvested 8 hours after KCl induction. PRPs protein expression levels were determined by immunoblotting. (A) Representative immunoblotting image of GluA1, GluA2, and PSD-95 protein expression. GAPDH is used as a loading control. (B) Quantification of PRPs protein expression. GluA1 expression is upregulated in the presence of ASO combined with neuronal stimulation. Treatment with ASO leads to an increase of PSD-95 protein level in primary cortical neurons (one-way ANOVA with Sidak’s *post hoc* tests. \**P* < 0.05, *n*.*s*. not significant. (C) Representative images of immunoblotting analysis showing NR2B protein expression. GAPDH is used as a loading control. (D) Quantification of NR2B protein expression. ASO treatment induces an increase in NR2B expression (one-way ANOVA with Sidak’s *post hoc* tests. \**P* < 0.05). Data are presented as mean ± SEM. The analysis revealed that the steady-state levels of GluA1 are elevated when the CPEB3 ribozyme is inhibited, but these levels do not increase further upon KCl stimulation of the cultured cortical neurons.

**S3 Fig.**
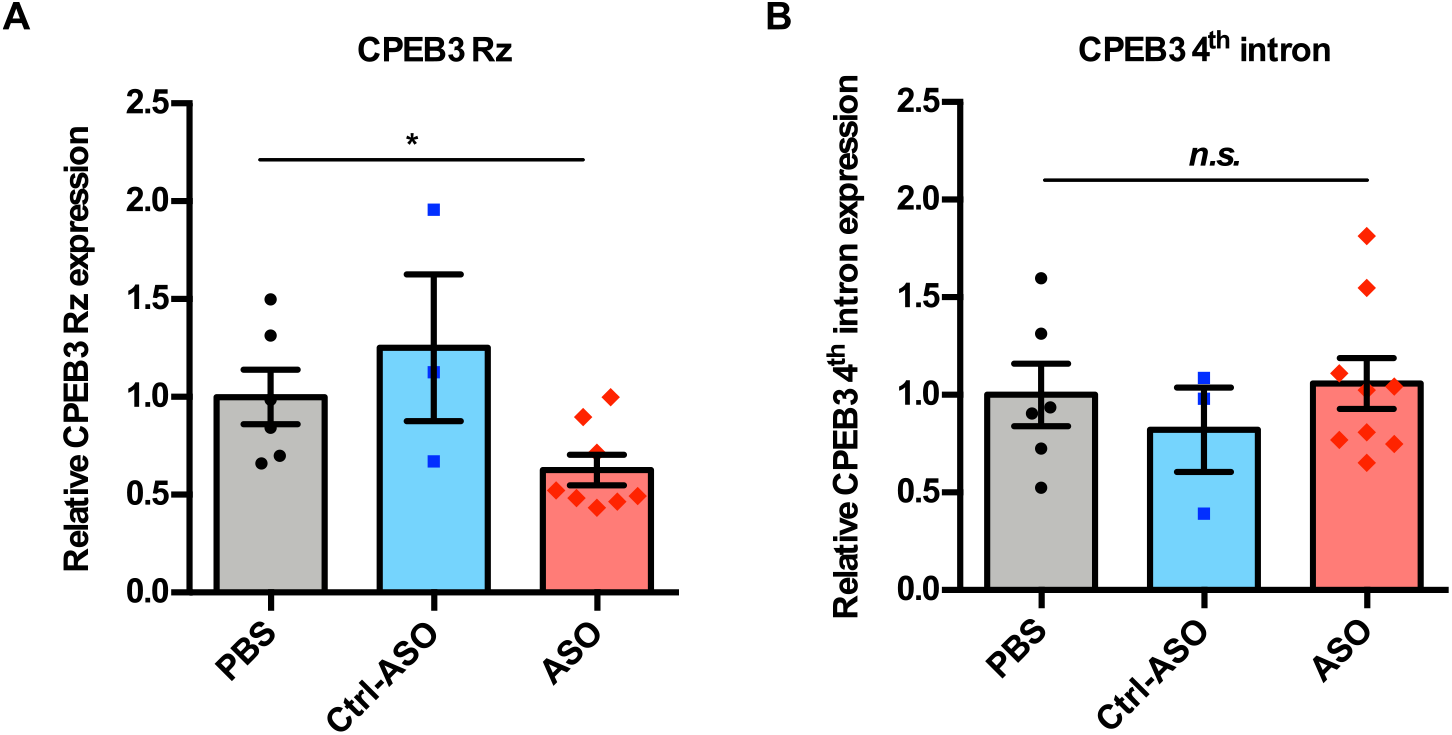
Inhibition of CPEB3 ribozyme by ASO *in vivo*. (A) Validation of CPEB3 ribozyme knockdown *in vivo*. Administration of CPEB3 ribozyme ASO to the mouse CA1 hippocampus leads to a decrease in CPEB3 ribozyme levels (one-way ANOVA with Sidak’s *post hoc* tests \**P* < 0.05). (B) The ribozyme ASO has high specificity for its cleavage site (in the 3^rd^ intron) *in vivo*. qRT-PCR analysis of the 4^th^ intron of *CPEB3* gene demonstrates no significant difference between controls and ASO groups (one-way ANOVA with Sidak’s *post hoc* test, *n*.*s*. not significant). Data are presented as mean ± SEM.

**S4 Fig.**
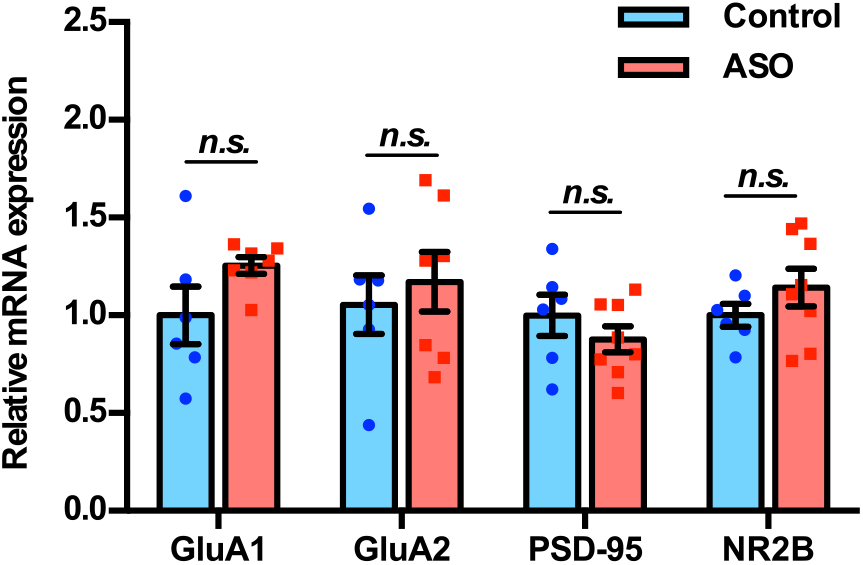
Inhibition of CPEB3 ribozyme does not affect transcription of other plasticity-related genes. qRT-PCR analysis of mature GluA1, GluA2, PSD-95, and NR2B mRNAs. No significant difference between ASO and control was observed for splice junctions within the mRNAs, showing that modulation of the CPEB3 ribozyme does not affect transcription or splicing of these mRNAs (unpaired *t* test, *n*.*s*. not significant). Data are presented as mean ± SEM.

**S5 Fig.**
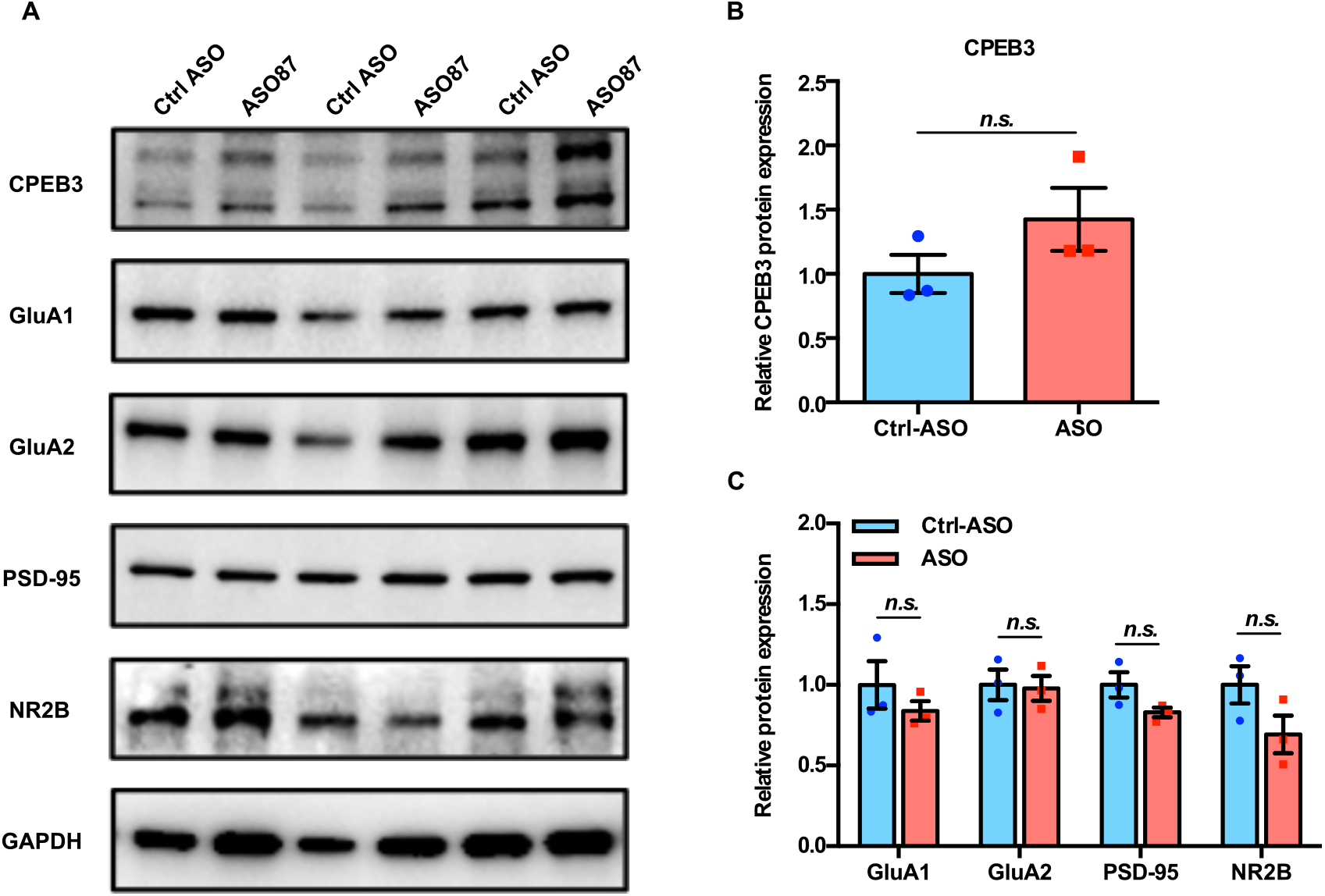
Effect of CPEB3 ribozyme on overall protein expression in the dorsal hippocampus. (A) Representative images of immunoblotting analysis. GAPDH is used as a loading control. (B) Quantification of CPEB3 protein expression (unpaired *t* test, *n*.*s*. not significant). (C) Quantification of PRPs protein expression (unpaired *t* test, *n*.*s*. not significant). Data are presented as mean ± SEM.

**S1 Table.** Kinetic parameters of murine CPEB3 ribozyme constructs^2^.

| Construct<sup>1</sup> | A | $k_1$ | B | $k_2$ | C |
| --- | --- | --- | --- | --- | --- |
| -10/72 | $0.72 \pm 0.09$ | $0.39 \pm 0.09$ | | | $0.082 \pm 0.026$ |
| -49/72/165 | $0.88 \pm 0.02$ | $0.42 \pm 0.04$ | $0.013 \pm 0.015$ | $0.11 \pm 0.03$ | $0.04 \pm 0.02$ |
| -233/72/165 | $0.78 \pm 0.04$ | $0.31 \pm 0.04$ | $0.035 \pm 0.006$ | $0.17 \pm 0.02$ | $0.029 \pm 0.005$ |

<sup>1</sup> Construct size is defined as the length of sequence upstream of the ribozyme cleavage site/CPEB3 ribozyme (72 nts)/downstream sequence.
<sup>2</sup> Co-transcriptional self-scission was modeled by a bi-exponential decay model with a residual. A and B represent fractions of the population cleaving with fast ( $k_1$ ) and slow ( $k_2$ ) rate constants, respectively. The residual (C) is interpreted as a fraction of the population that does not self-cleave. Errors represent SEM of at least three experiments. For the smallest ribozyme construct (-10/72), a monoexponential decay function was sufficient to model the data.

**S2 Table.**
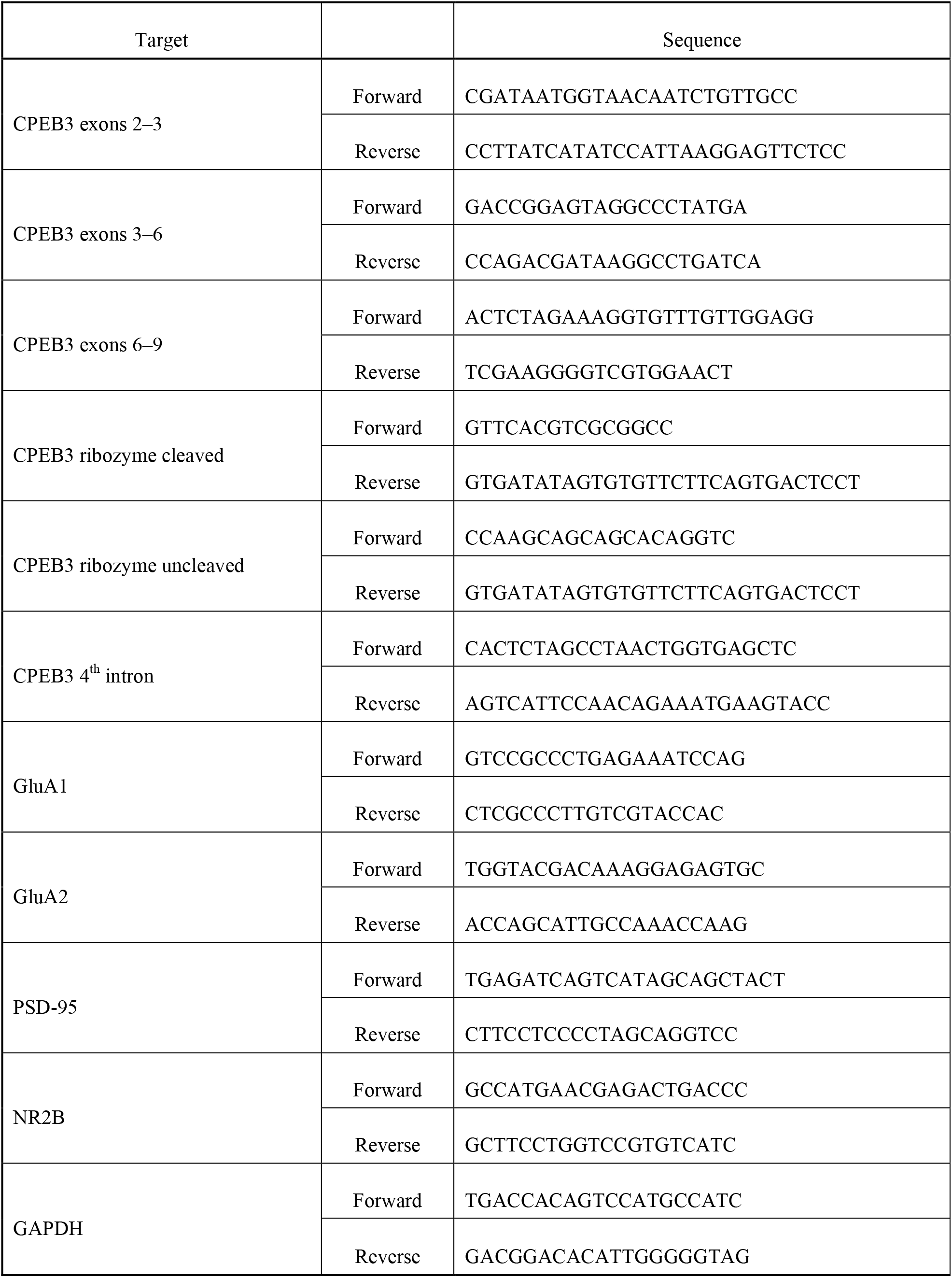
Primers used in qPCR.

**S3 Table.** Antibodies used in immunoblotting analysis.

| Antigen | Species | Company | Catalog # | Dilution |
| --- | --- | --- | --- | --- |
| CPEB3 | Rabbit | Abcam | ab18833 | 1:1,000 |
| GluA1 | Mouse | UC Davis/NIH NeuroMab Facility | 75-327 | 1:1,000 |
| GluA2 | Rabbit | Proteintech | 11994-1-AP | 1:2,000 |
| PSD-95 | Rabbit | Proteintech | 20665-1-AP | 1:2,000 |
| NR2B | Rabbit | Proteintech | 21920-1-AP | 1:2,000 |
| GAPDH | Mouse | Proteintech | 60004-1-Ig | 1:10,000 |
| Anti-rabbit HRP | Donkey | Thermo Fisher Scientific | A16023 | 1:10,000 |
| Anti-mouse HRP | Goat | R&D system | HAF007 | 1:1,000 |

**S4 Table.**
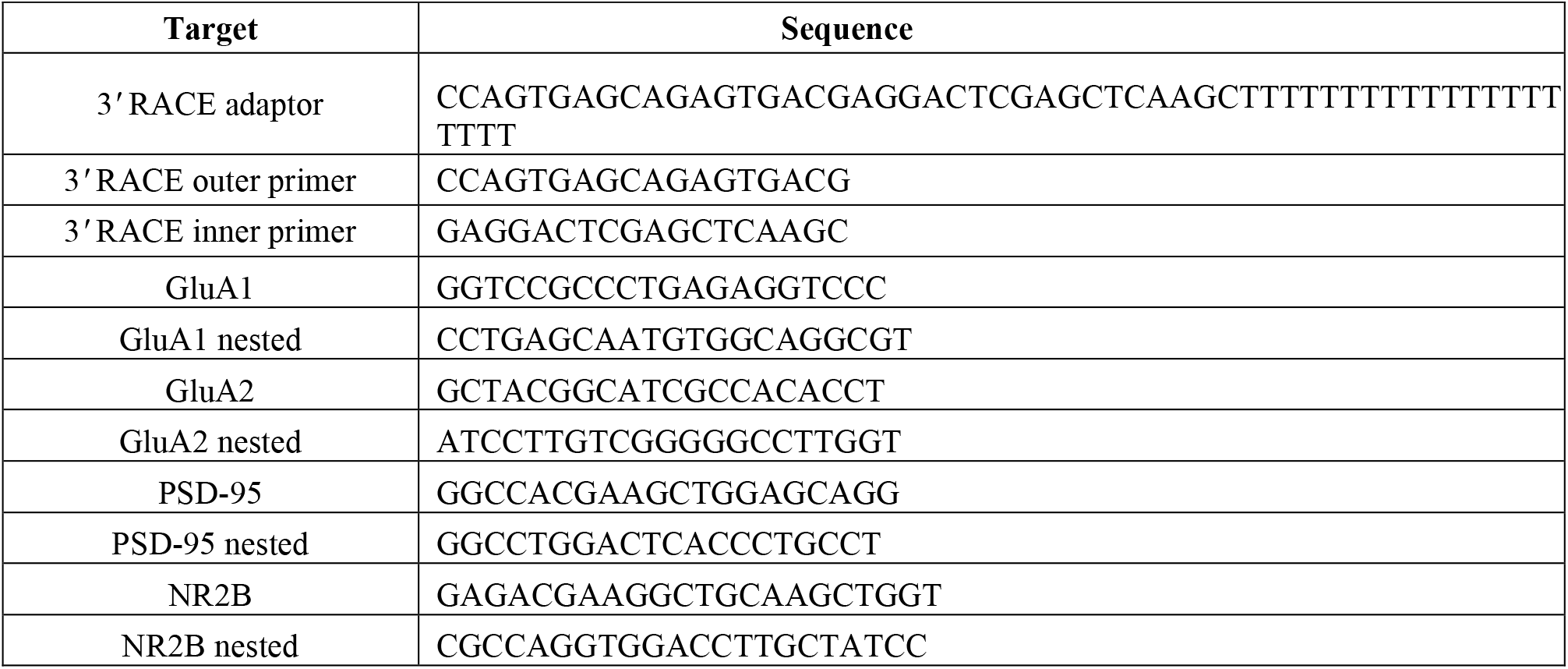
Primers used in 3’ RACE.

## Notes

### Competing Interest Statement

The authors have declared no competing interest.

### Summary of Updates

In this version we added references and reorganized the Introduction and Discussion sections.

